# Single-nucleus full-length RNA profiling in plants incorporates isoform information to facilitate cell type identification

**DOI:** 10.1101/2020.11.25.397919

**Authors:** Yanping Long, Zhijian Liu, Jinbu Jia, Weipeng Mo, Liang Fang, Dongdong Lu, Bo Liu, Hong Zhang, Wei Chen, Jixian Zhai

**Author notes:** These authors contributed equally to this work (Yanping Long, Zhijian Liu, Jinbu Jia). Correspondence (J.Z.).

## Abstract

The broad application of large-scale single-cell RNA profiling in plants has been restricted by the prerequisite of protoplasting. We recently found that the Arabidopsis nucleus contains abundant polyadenylated mRNAs, many of which are incompletely spliced. To capture the isoform information, we combined 10x Genomics and Nanopore long-read sequencing to develop a protoplasting-free full-length single-nucleus RNA profiling method in plants. Our results demonstrated using Arabidopsis root that nuclear mRNAs faithfully retain cell identity information, and single-molecule full-length RNA sequencing could further improve cell type identification by revealing splicing status and alternative polyadenylation at single-cell level.

## Background

High-throughput single-cell transcriptome studies have thrived in animal and human research in recent years[1-5]. However, despite successful single-cell characterization at a relatively low scale in maize developing germ cells[6] and rice mesophyll cells[7] using capillary-based approaches[8], only a handful of large-scale single-cell RNA studies using high-throughput platforms such as 10x Genomics or Drop-seq[9] have been published in plants[10], all of which profiled protoplasts generated from the root of Arabidopsis[11-17]. A major reason for this narrow focus of tissue type is that plant cells are naturally confined by cell walls, and protoplasting is required to release individual cells – a procedure that is thoroughly tested for Arabidopsis roots[18-20] but remains to be difficult or impractical in many other tissues or species. Moreover, generating protoplasts from all cells uniformly is challenging given the complexity of plant tissues, and the enzymatic digestion and subsequent cleanup process during protoplast isolation may trigger the stress response and influence the transcriptome. Therefore, a protoplasting-free method is urgently needed to broaden the application of large-scale single-cell analysis in plants.

We recently characterized full-length nascent RNAs in Arabidopsis and unexpectedly found a large number of polyadenylated mRNAs that are tightly associated with chromatin[21]. Since it is considerably easier and more widely applicable to perform nuclei isolation on various plant tissues than protoplasting, we set out to test if the polyadenylated RNAs in a single nucleus are sufficient to convey information on cell identity using the 10x Genomics high-throughput single-cell platform. Besides the standard Illumina short-read library which primarily captures abundance information, long-read sequencing has recently been incorporated into single-cell studies [22-24]. To access the large number of intron-containing RNAs in plant nuclei, we also constructed a Nanopore-based long-read library and developed a bioinformatic pipeline named “snuupy” (<u>s</u>ingle <u>nu</u>cleus <u>u</u>tility in <u>p</u>ython) to characterize mRNA isoforms in each nucleus (Figure 1a, Supplemental Figure 1). This long-read single-nucleus strategy would enable plant biologists to bypass protoplasting, study RNA isoforms derived from alternative splicing and alternative polyadenylation (APA) at the single-cell level, and provides additional dimensions of transcriptome complexity that could potentially further improve clustering of different cell types.

**Figure 1.**
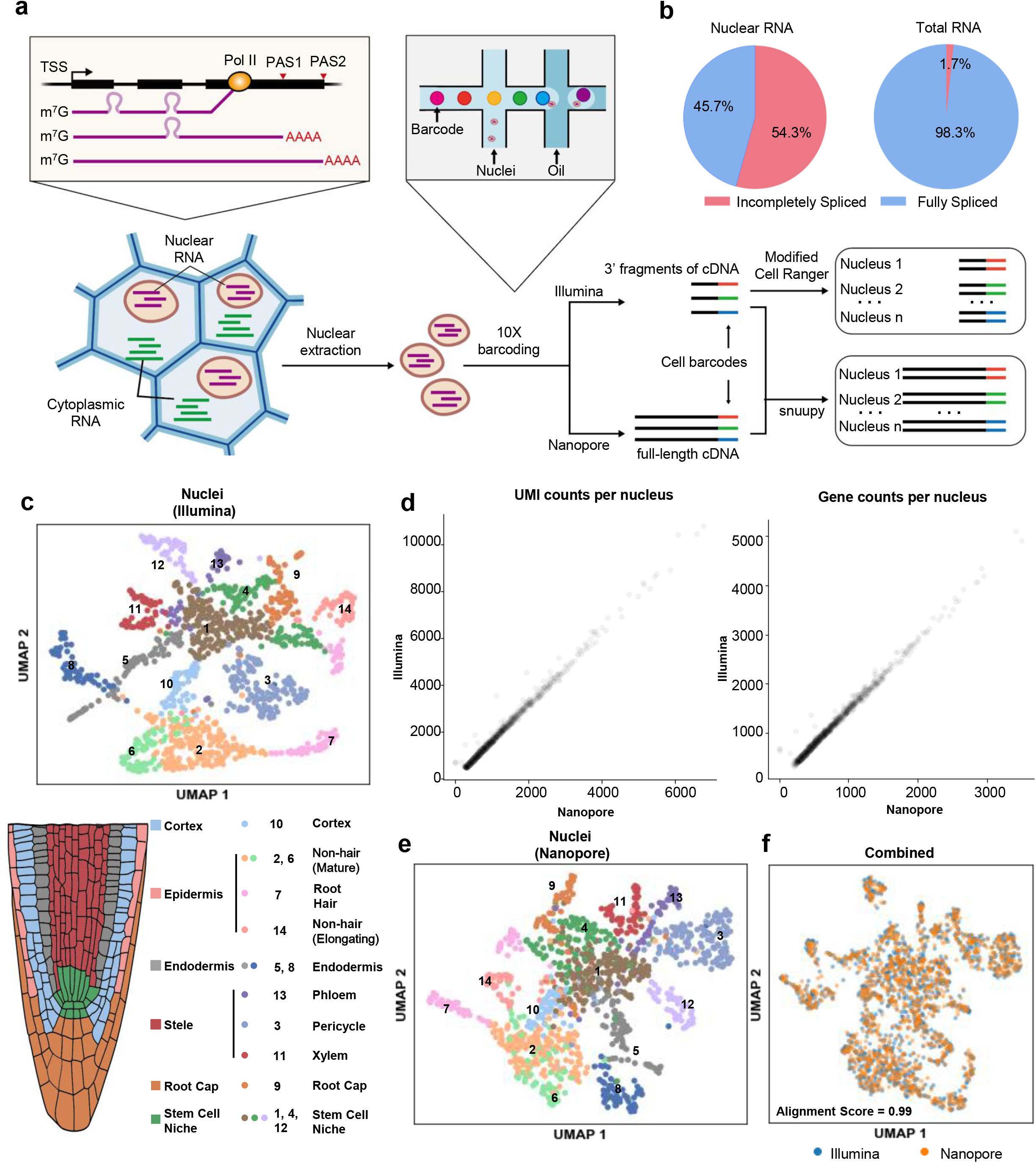
Protoplasting-free large-scale single-nucleus RNA-seq reveals the diverse cell types in Arabidopsis root. **a**, Schematic diagram of protoplasting-free single-nucleus RNA-seq. **b**, Incompletely spliced and fully spliced fractions of the Nanopore reads from our single-nucleus RNA library, compared with a previously published total RNA nanopore library[26]. **c**, UMAP visualization of the root cell types clustered using Illumina single-nucleus data (upper panel), and cartoon illustration of major cell types in Arabidopsis root tip (lower panel). **d**, Numbers of UMIs (left) and genes (right) detected in each nucleus from the Illumina and Nanopore data. **e**, UMAP visualization of the root cell types clustered using abundance information from the Nanopore single-nucleus data. The cell color is the same as in Figure 1c. **f**, UMAP visualization of the integration of two datasets. The batch effect is removed by Scanorama. Alignment score is calculated by Scanorama[27] and in the range from 0 to 1. Higher alignment score indicates higher similarity between a pair of datasets.

## Results and discussion

Here, we chose to use the Arabidopsis root to validate the effectiveness of our protoplasting-free single-nuclei RNA sequencing approach because of the well-studied cell types[25] and the rich resource of single-cell data[11-16] of this tissue. We directly isolated nuclei by sorting from homogenized root tips of 10-day-old Arabidopsis seedlings without protoplasting (Supplemental Figure 2). The nuclei were fed to the 10x Genomics Chromium platform to obtain full-length cDNA templates labeled with nucleus-specific barcodes, which are subsequently divided into two equal parts and used for constructing Illumina short-read and Nanopore long-read libraries, respectively (Figure 1a).

From the Illumina library, we obtained a total of 1,186 single-nucleus transcriptomes covering 18,913 genes, with median genes/nucleus at 810 and median UMIs/nucleus at 1131. It is worth noting that the proportion of intron-containing mRNAs is extremely high in plant nucleus - 54% compared to less than 2% in total RNAs[26] (Figure 1b). After generating the cell-gene abundance matrix from Illumina data, we used the Scanorama algorithm[27] to compare our dataset with several recently published root single-cell datasets from protoplasts[11, 12, 14-16]. The expression abundance matrix from our single-nucleus dataset closely resembles the protoplasting-based single-cell dataset generated from the same tissue (10-days seedling, 0.5 mm primary root tips)[15] (Supplemental Figure 3). Next, we utilized an unbiased graph-based clustering method Louvain[28] and identified 14 distinct cell clusters (Figure 1c). We then applied a set of cell type-specific marker genes provided in a recent massive single-cell study of Arabidopsis roots[17] to annotate each cluster (See Methods, Supplemental Table 1). We were able to assign cell types to all 14 clusters and identified 10 major root cell types previously reported (Figure 1c, Supplemental Figure 4), with the signature transcripts for each cell type enriched in the corresponding cluster (Supplemental Figure 5, Supplemental Figure 6). Consistent with previous reports[11-16], we also noticed that some cell types from our result are composed of multiple clusters, such as Stem Cell Niche (cluster 1, 4 and 12), mature Non-hair (cluster 2 and 6), Endodermis (cluster 5 and 8) (Figure 1c), demonstrating additional heterogeneity (subcell types) within cell types. Moreover, we found the exact same subcell type marker genes of endodermis are enriched in each of its corresponding subcell types as shown in Zhang *et al.* [15] (Supplemental Figure 7), demonstrating the robustness of our single-nucleus data. Taken together, we demonstrated that transcriptomes of single nucleus are sufficient for cell type identification, and can be used as a reliable alternative to protoplasts.

As to the Nanopore data analysis, a key challenge is that the relatively low sequencing accuracy of Nanopore (~95% per base) makes it difficult to correctly recognize the cell barcodes and UMI information on each Nanopore read. To solve this problem, Lebrigand *et al.* developed a method named Sicelore to use Illumina short reads generated from the same cDNA library as the guide to allocate Nanopore reads [22]. Sicelore searches for both polyA and adapter sequence and define the region between these two as the potential barcode and UMI. However, this algorithm relies on the recognition of polyA tail sequence generated by the Nanopore basecalling software, which tends to severely underestimate the length of polyA tail [29]. We tried to further improve Sicelore by developing a polyA independent algorithm (named snuupy), which searches for cell barcodes and UMIs in the unmapped region of Nanopore reads (See Methods and Supplemental Figure 1, Supplemental Figure 8a). As the result, snuupy recovers 20% more reads from our Nanopore data compared to using Sicelore [22] (Supplemental Figure 8b). After snuupy processing, we obtained 1,169 long-read single-nucleus transcriptomes from Nanopore data (compared to the 1,186 from Illumina data). The median UMI counts per nucleus (729) and the median gene counts per nucleus (563) from Nanopore data are ~64% and ~70% of the Illumina count, respectively, and highly consistent in all nuclei (Figure 1d). The clustering result using Nanopore abundance matrix closely resembles the one generated by Illumina data (Figure 1e, Figure 1f), suggesting that Nanopore data itself is sufficient for cell-type classification, consistent with a recent large-scale single-cell analysis in human and mouse cells performed entirely with Nanopore data[22, 23].

The single-nucleus long-read Nanopore library provides isoform-level information such as splicing and APA, compared to Illumina library which only captures abundance information of transcripts. Therefore, we generated two additional isoform matrices to track splicing and APA in single nucleus, respectively (Figure 2a and Supplemental Figure 9), and combined them with the Illumina abundance matrix for a multilayer clustering, to test if these extra layers of information could improve cell type classification. Indeed, we found that the original cluster 2 (Mature Non-hair) and cluster 10 (Cortex) from Illumina data (Figure 1c) can be further separated into two subcell type clusters after the multilayer clustering (Figure 2a). As an example, from the Illumina data, transcripts of AT3G19010 are present in both subcell type 2.1 and 2.2 (Figure 2b and 2c), while the Nanopore data revealed a large difference at the splicing level of this gene between the two sub-clusters, with the second intron largely unspliced in subcell type 2.2 (Figure 2d). It is worth noting that, *JAZ7*, the top1 enriched gene in cluster 2.2 (Figure 2e), can regulate splicing during jasmonate response [30], implying a fascinating potential of cell-type specific regulation of splicing that could be investigated in the future.

**Figure 2.**
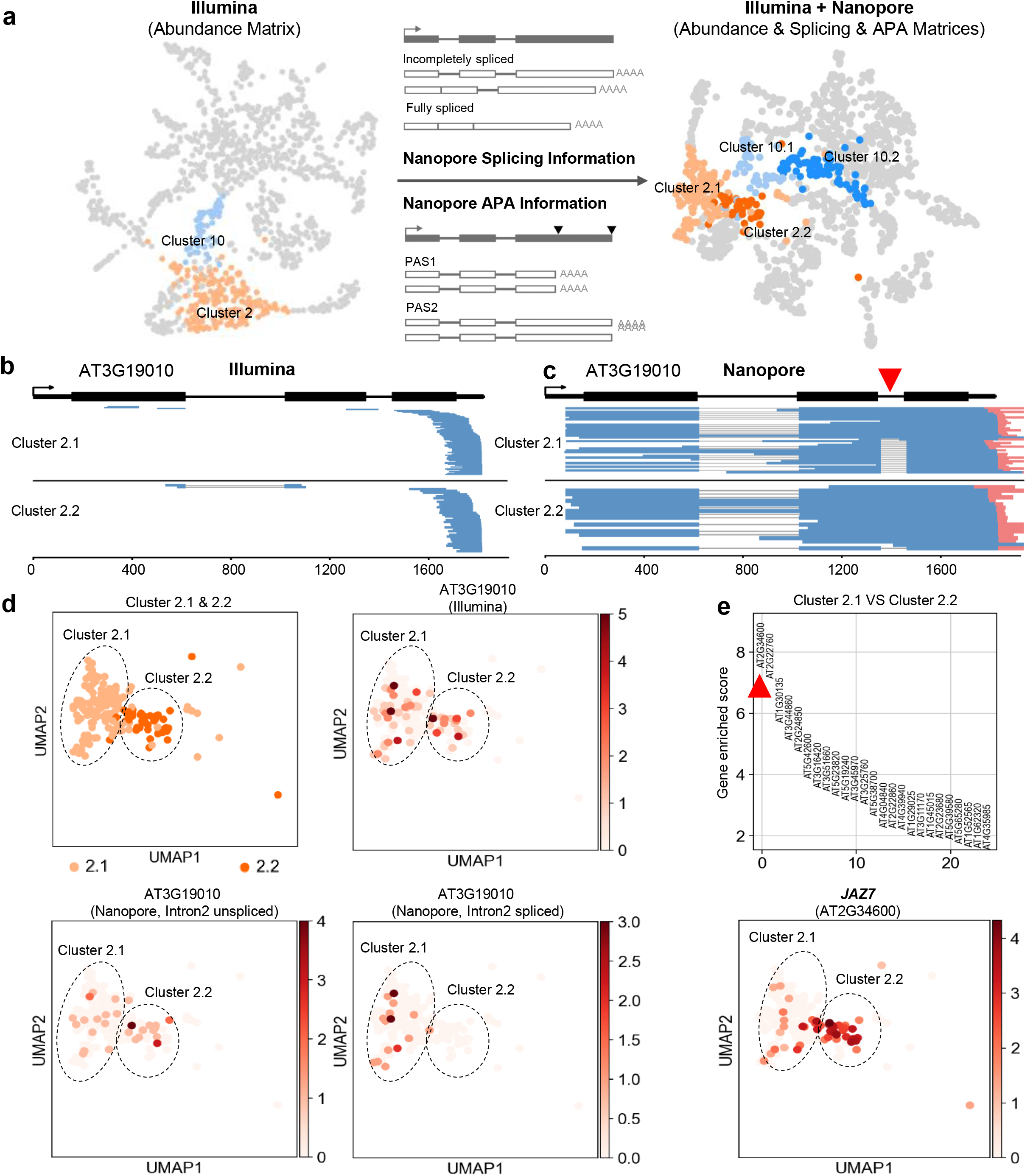
Nanopore long read single-nucleus RNA-seq improves cell type identification. **a**, Multi-layer matrices combining Illumina abundance matrix with Nanopore splicing and APA information improve cell type identification. **b,c**, Genome-browser plot of Illumina reads(**b**) and Nanopore reads(**c**) aligned to gene AT3G19010. The second intron of AT3G1910 shows different splicing patterns between Cluster 2.1 and Cluster 2.2. The red arrowhead indicates the second intron. Red bar at the 3’ end of Nanopore reads (blue) indicates the Poly(A) tail. **d**, UMAP visualization shows the abundance distribution of AT3G19010 as well as the differential splicing of the second intron between Cluster 2.1 and Cluster 2.2. **e**, The top 25 genes enriched in Cluster 2.2 are ranked by enriched score compared to Cluster 2.1 (upper panel) and UMAP visualization shows the abundance distribution of the most enriched gene *JAZ7* (lower panel). The enriched score is calculated using *rank_genes_groups function* of Scanpy. The red arrowhead indicates the most enriched gene in Cluster 2.2.

## Conclusions

According to previous reports in the animal system, especially for neurons and frozen materials, single nucleus generates comparable RNA to single cell and establishes a robust transcriptome atlas[31-34]. As a proof-of-concept demonstration in plant, our results showed that protoplasting-free large-scale single-nucleus sequencing is sufficient for cell type classification and marker gene identification in Arabidopsis roots. As we are preparing this manuscript, several groups have also recently adopted the nuclei-based protoplasting-free strategy independently to investigate various plant tissues[35-39]. Eliminating protoplasting as a prerequisite would enable large-scale single-cell profiling on a wide range of tissues and plant species. Our method uniquely combined Nanopore-based full-length RNA sequencing method with single-nuclei sequencing to capture isoform diversity at single-nucleus level, which can facilitate cell type classification by providing extra layers of information in addition to abundance.

## Methods

### Nuclei isolation from root tip of Arabidopsis

The wild-type Arabidopsis seedlings (Col-0) were grown on 1/2 MS plates at 22 °C (16 h light/8 h dark) for 10 days before harvest. The root tip region (5 mm) of seedlings were cut and transferred immediately into a 1.5 ml RNase-free Eppendorf tube kept in liquid nitrogen and were ground into fine powder by a 1000 μl pipette tip in the tube. The powder was then dissolved in 300 μl ice-cold Extraction Buffer (EB) - 0.4 M sucrose, 10 mM Tris-HCl pH 8.0, 10 mM MgCl_2_, 0.2% (w/v) Triton X-100, 1 mM dithiothreitol (DTT), 1× protease inhibitor (Roche), 0.4 U/μl RNase inhibitor (RNaseOUT, Thermo Fisher Scientific). Nonionic surfactant Triton X-100 is used to release nuclei, and avoid aggregation during FACS[40]. After gentle vertexing and inversion, the homogenate was filtered through a 20 μm cell strainer into a new tube. Another 400 μl EB was added to the strainer to wash the remaining nuclei. After centrifugation at 4 °C, 2000 g for 5 min, the supernatant was removed carefully to avoid RNA contaminants from the cytoplasmic fraction. The pellet was washed twice at 4 °C, 2000 g, 5 min with 1 ml EB, and then resuspended in 500 μl EB. For sorting, the nuclei were stained with 4,6-Diamidino-2-phenylindole (DAPI) and loaded into a flow cytometer with a 70 μm nozzle. 1 ml EB was used as the collection buffer. A total of 40,000 nuclei were sorted based on the DAPI signal and the nuclear size. To avoid aggregation, the sorted nuclei were pelleted at 4 °C, 2000 g, 5 min, and then resuspended in 100 μl PBST buffer (1× PBS with a low concentration of 0.025% Triton X-100). After checking the quality of nuclei and counting under a microscope using the DAPI channel, 5000 nuclei were transferred into a new tube with 500 μl PBST buffer and centrifuged at 4 °C, 2000 g, 5 min. Then the pellet was resuspended in 20 μl PBST buffer.

### Single nucleus RNA-seq library construction for Illumina and Nanopore sequencing

Libraries were constructed according to the standard 10x Genomics protocol (Single Cell 3’ Reagent Kits v2 User Guide) with modifications to accommodate Nanopore long-read sequencing. Briefly, nuclei suspension from the previous step (~5000 nuclei) were loaded onto the 10x Genomics ChIP, and libraries were made using a 10x Chromium Single Cell 3’ Solution V2 kit. To obtain full-length cDNA, we extend the elongation time during cDNA amplification from the standard 1 min to 2 minutes. Half of the cDNA template was used to construct Illumina library according to the manufacturer’s instruction and sequenced with Illumina NavoSeq (Read1:28 bases + Read2:150 bases); the other half of the template was used to make Nanopore library using the Oxford Nanopore LSK-109 kit and sequenced on a MinION flow cell (R9.4.1).

### Illumina single-nuclei data analysis

Raw reads were mapped to the TAIR10 reference genome by Cell Ranger (v3.1.0) using the default parameters. Cell Ranger (v3.1.0) only counts reads without introns; to accommodate the high proportion of intron-containing reads in our single-nucleus libraries, we removed the intron regions of each read and re-aligned reads to the reference genome by Cell Ranger to identify the nuclei barcode, UMI, and corresponding gene of each read (Supplemental Figure 1). For quality control purpose, genes expressed in less than three nuclei were discarded, and cells with gene counts more than 2300 or fewer than 350 were removed. The Illumina abundance matrix was subsequently analyzed using Scanpy package (v1.6.0)[41] with recommended parameters for normalization, log-transformation, and scaling. Then principal component analysis and Louvain algorithm were used on this abundance matrix for clustering. Next, we used the marker genes for different cell types identified in a massive single-cell root data [17] (Supplemental Table 1) to annotate the cell type of each cluster. We first calculate the cell score of each cell type for all cells based on the enrichment degree of a given marker gene set in a given cell, as previously described method [42]. If the highest score exceeds zero, the cell is assigned to the corresponding cell type; otherwise it is assigned as unknown (Supplemental Figure 4a). Then each cluster was annotated as the cell type with the highest proportion (Supplemental Figure 4b), and we used developmental stage specific genes identified in the massive single-cell root data [17] (Supplemental Table 1) to further annotate the clusters resenting non-hair cells as either mature non-hair or elongating non-hair cells (Supplemental Figure 4c).

Five previously published single-cell RNA-seq data of protoplasted Arabidopsis roots using 10x Genomics platform were collected from public databases[11, 12, 14-16]. We use Scanorama[27] to remove batch effects and calculate the alignment score between different datasets.

### Nanopore single-nuclei data processing and isoform analysis

Raw Nanopore data were basecalled using Guppy (v3.6.0) with the parameters “--c dna_r9.4.1_450bps_hac.cfg --fast5_out”. The basecalled reads were mapped to the TAIR10 genome by minimap2 (v2.17) with the parameters “-ax splice --secondary=no-uf --MD --sam-hit-only”, and the multi-mapped reads as well as potential chimeric reads (either the 5’ or 3’ unmapped region is great than 150 nt) were filtered out. The nucleus barcodes and UMI sequences in Nanopore reads were extracted from the unmapped sequences of each read via aligning against all barcode/UMI combinations identified from the Illumina library made from the same full-length cDNA templates, a strategy inspired by the algorithm Sicelore[22]. To reduce search space, we divided the genome into non-overlapped 500-bp bins, and only matched the Illumina barcode/UMI combinations from the bins overlapping or adjacent to the mapping genome region of specific Nanopore read (Supplemental Figure 6). To speed up the alignment process, we first used the heuristic algorithm Blastn (v2.10.0) to find potential seed regions with parameters “-word_size 7-gapopen 0 -gapextend 2-penalty-1-reward 1” and then re-aligned the seed regions by the more accurate Smith-Waterman local alignment algorithm. Our pipeline assigns the closest barcode-UMI match (i.e. with minimal mismatch/gap) to each Nanopore read, allowing up to three base errors (mismatch/gap) for either barcode or UMI, and remove reads with multiple best matching barcode-UMIs. After the barcode and UMI assignment, the Nanopore reads with the same UMI were used to generate an error-corrected consensus sequence of the original RNA molecule by poaV2[43] and racon[44]. PAS isoform annotation and the intron splicing status of Nanopore read were determined as previously described[21, 45]. The resulted APA and splicing matrices for all nuclei were merged with Illumina abundance matrix and analyzed by Scanpy.

The same Cell Ranger result is used as the input file for Sicelore. Except that the maximum edit distance during Barcode and Umi assignment is forcibly set to 3, the remaining parameters are the same as the official example (https://github.com/ucagenomix/sicelore/blob/master/quickrun.sh).

### Data and software Availability

All data generated in this study were deposited in NCBI with accession PRJNA664874 (https://www.ncbi.nlm.nih.gov/bioproject/PRJNA664874). The snuupy package for single-nucleus Nanopore data processing can be accessed at https://github.com/ZhaiLab-SUSTech/snuupy.

## Acknowledgments

Group of J.Z. is supported by the National Key R&D Program of China Grant (2019YFA0903903), the Program for Guangdong Introducing Innovative and Entrepreneurial Teams (2016ZT06S172), the Shenzhen Sci-Tech Fund (KYTDPT20181011104005), and Key Laboratory of Molecular Design for Plant Cell Factory of Guangdong Higher Education Institutes (2019KSYS006).

## Author Contributions

Y.L., L.F., D.L. and B.L. performed the experiments. Y.L., Z.L., J.J., W.M. and H.Z. analyzed the data. J.Z., W.C. and J.J. oversaw the study. All authors wrote and revised the manuscript.

---

**Supplemental Figure 1.**
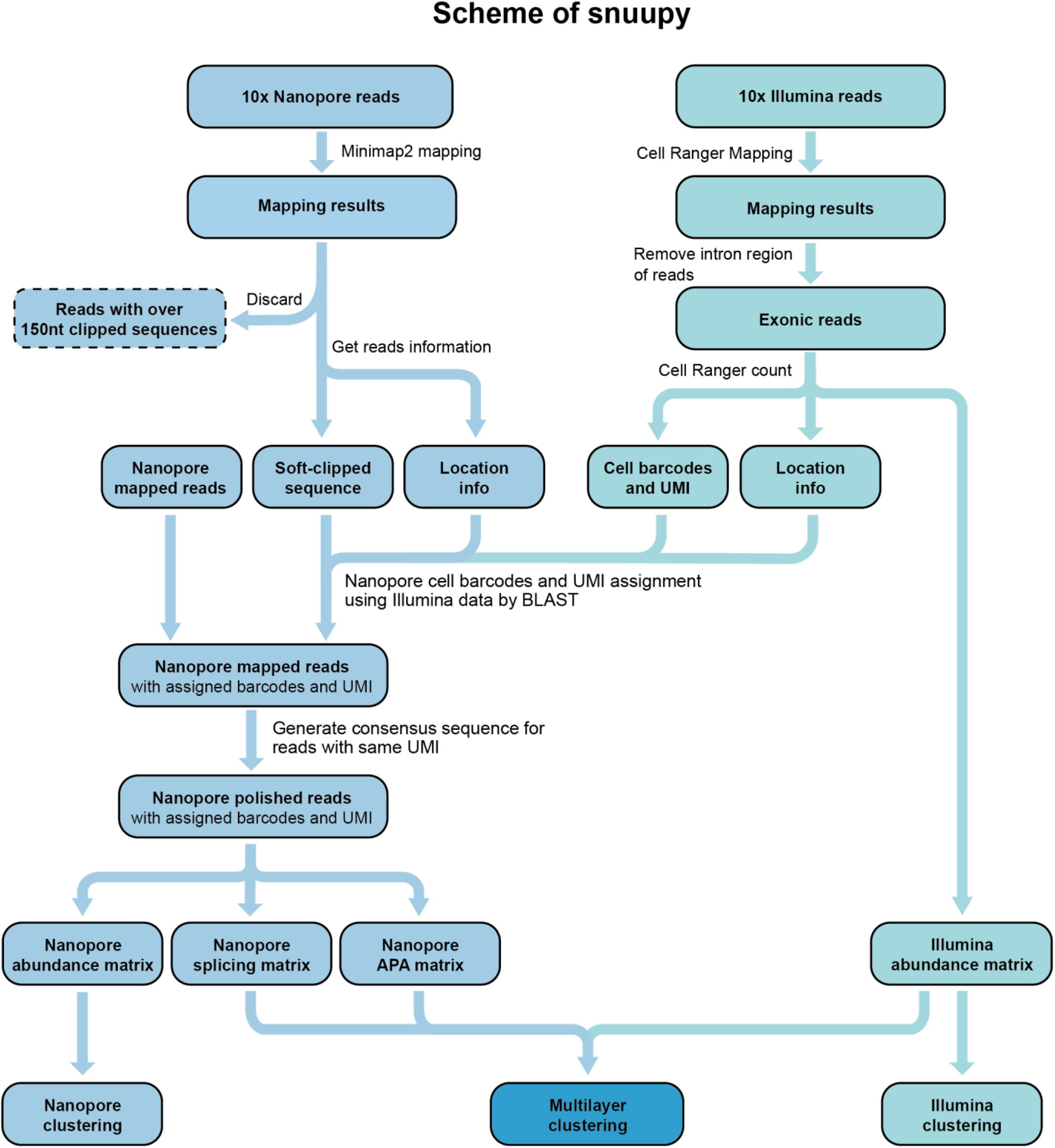
Schematic diagram of snuupy bioinformatic pipeline.

**Supplemental Figure 2.**
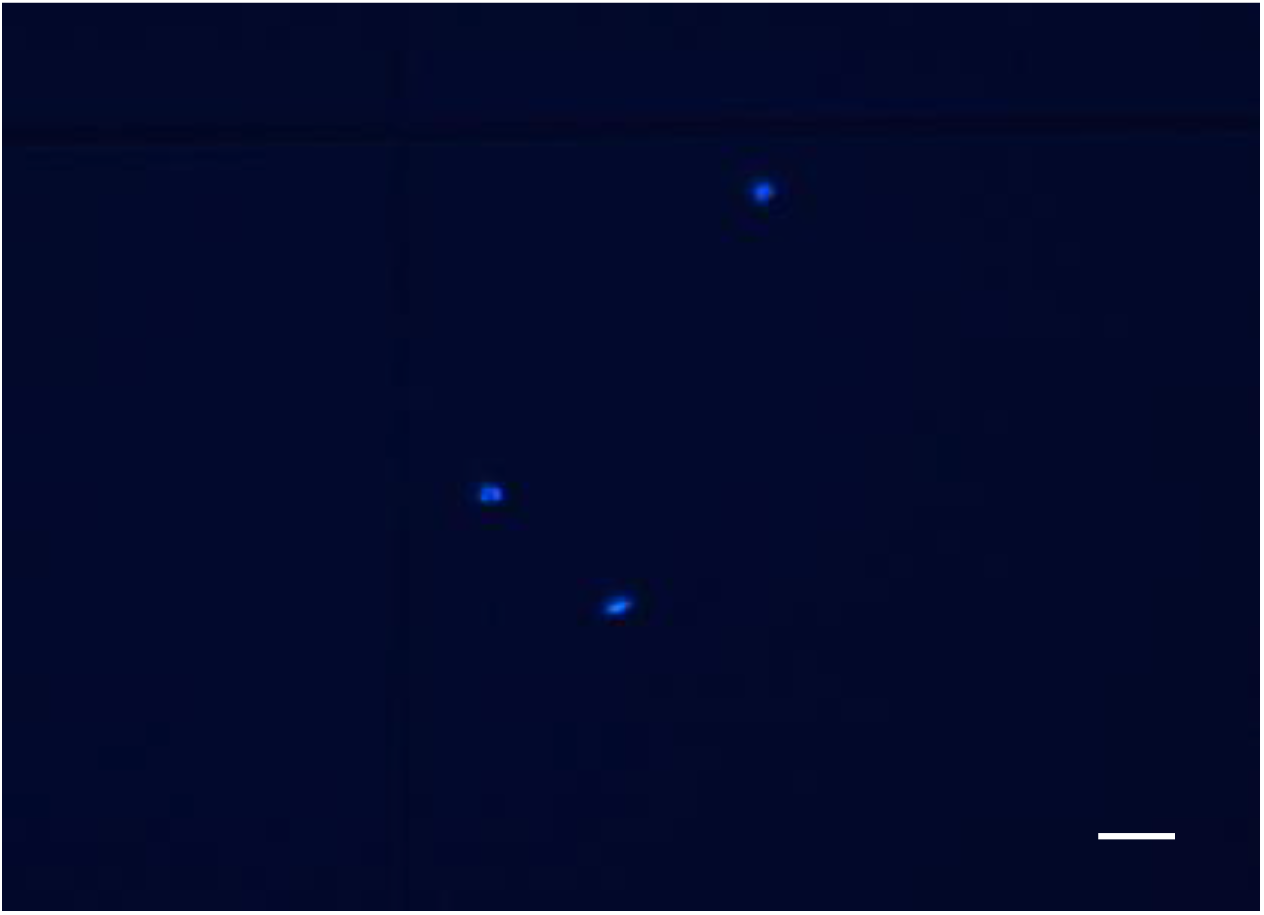
The sorted nuclei were observed under a microscopy with DAPI staining. Bar = 20 μm.

**Supplemental Figure 3.**
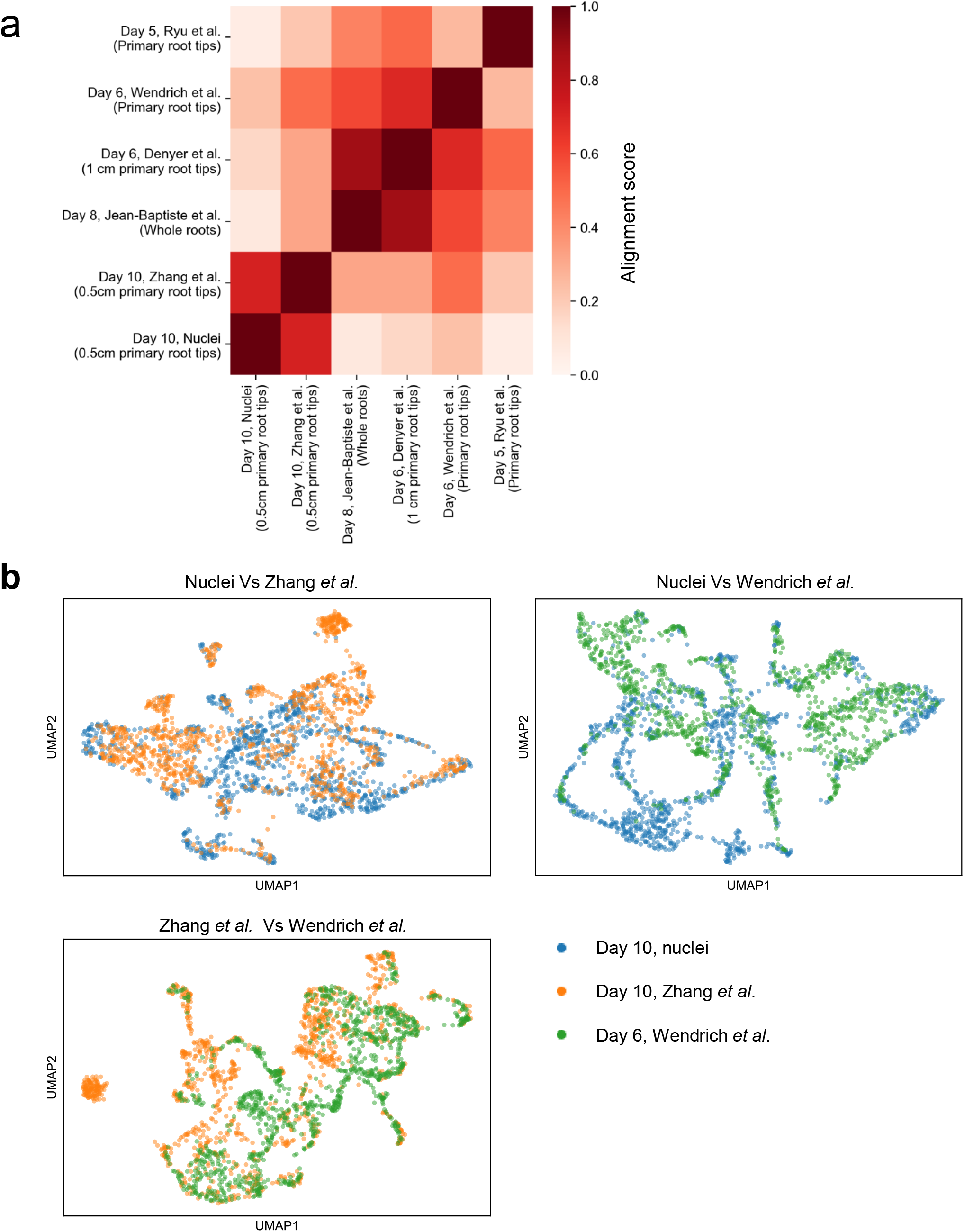
Dataset generated by snRNA-seq is consistent with protoplast-based scRNA-seq. **a**, Heatmap represents alignment score between the single-nucleus data and single-cell datasets generated from 10x Genomics platform. Alignment score is calculated by Scanorama[27]. Higher alignment score indicates higher similarity between a pair of datasets. **b**, Pairwise integration of three single cell/nucleus datasets. The batch effect is removed by Scanorama. The expression matrix is downsampled to the same dimension as the single-nucleus data.

**Supplemental Figure 4.**
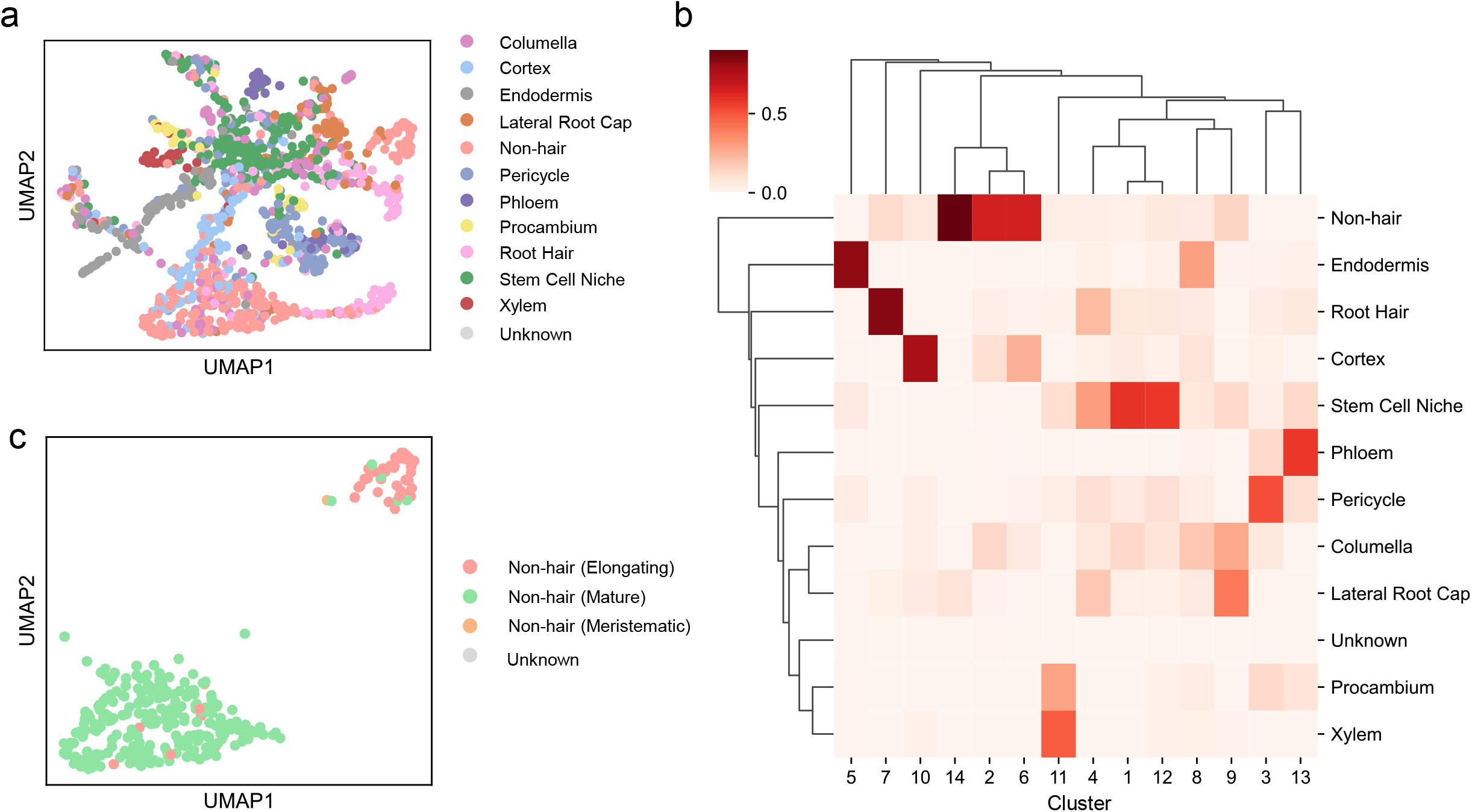
Identification of clusters by a marker-gene-based method. We calculate the cell score[41] for each cell based on type-specific genes[17]. Cells are classified as the type with the highest cell score. **a**, UMAP visualization of the 1186 cells. Colors denote corresponding cell types. **b**, Heatmap visualization of the proportion of cell types in each cluster. **c**, UMAP visualization of the cells within cluster 2, cluster 6, cluster 14. The developmental stage specific genes of non-hair cells are used to calculate the cell score and annotate each cell.

**Supplemental Figure 5.**
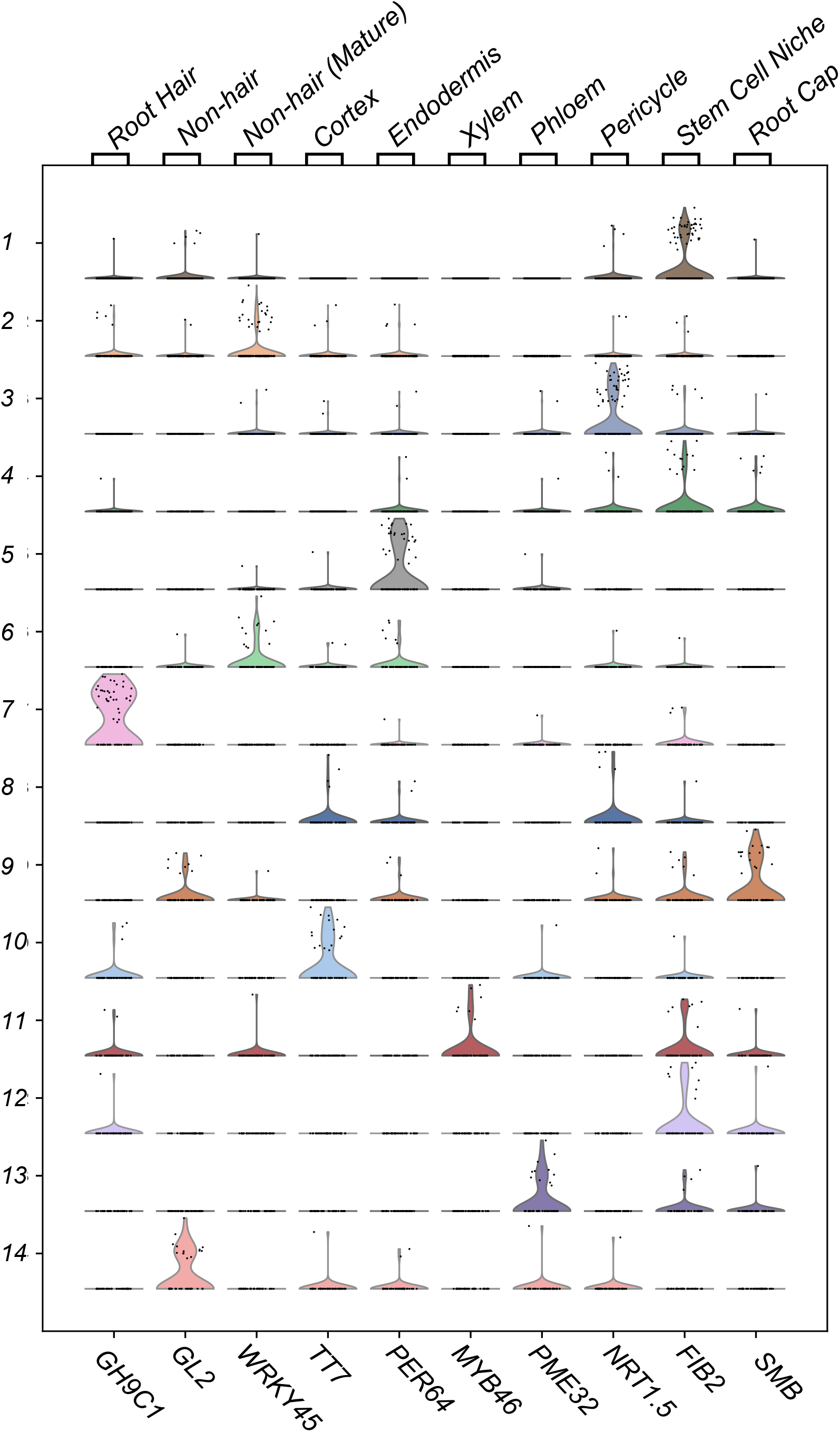
Violin plots showing the expression levels of previously reported cell type specific marker genes in 14 clusters.

**Supplemental Figure 6.**
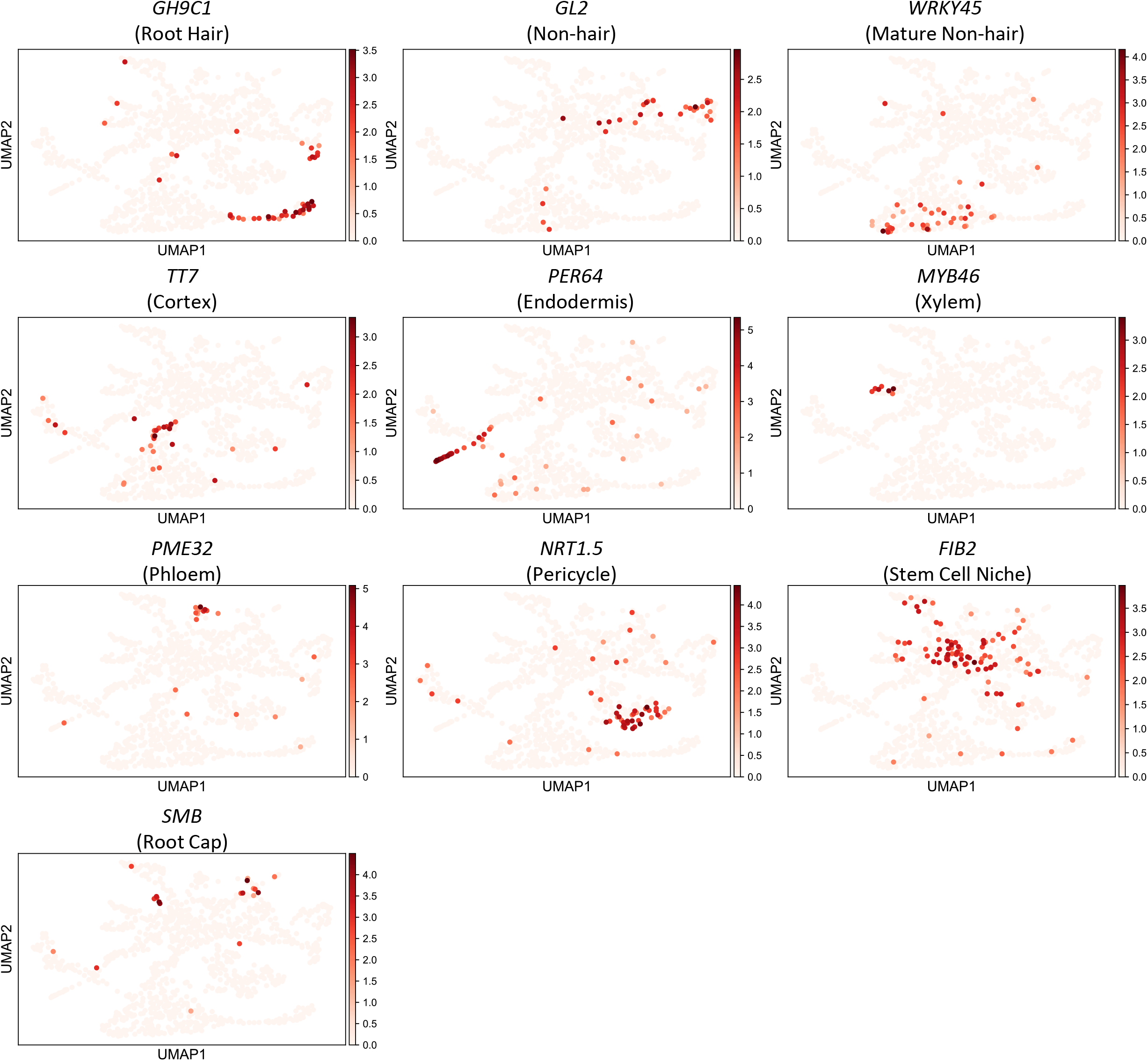
UMAP visualization of the representative cell-type marker genes for each of the 14 cell clusters. The cell clusters and UMAP visualization are the same as those shown in Figure 1c. Color intensity indicates the relative expression level.

**Supplemental Figure 7.**
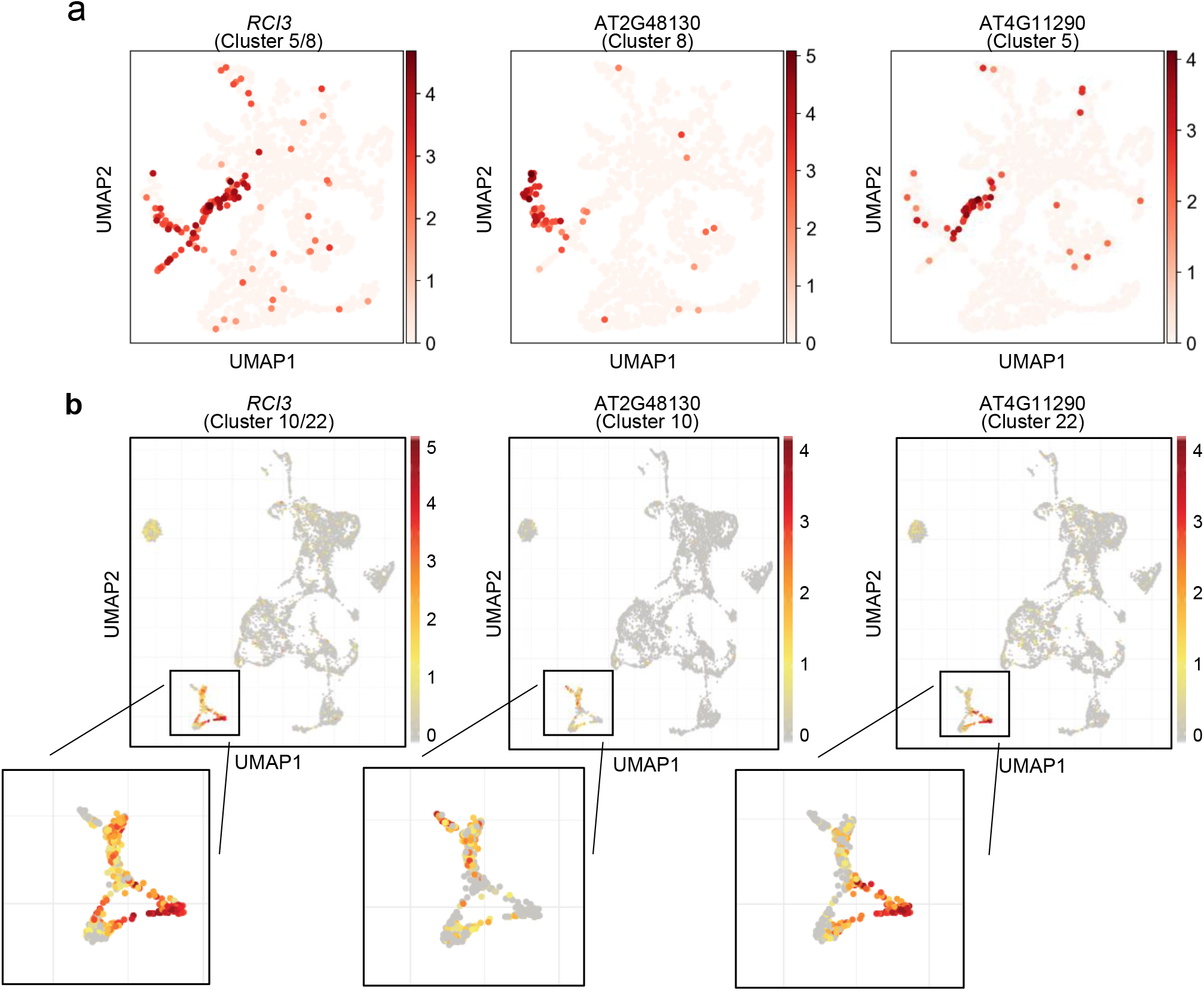
UMAP visualization showing the abundances of representative marker genes in two subcell types of endodermis. The protoplasting-free single-nucleus RNA-seq data (**a**) can also accurately identify the subtypes as previously published protoplasting-based single-cell RNA-seq data (**b**)[15].

**Supplemental Figure 8.**
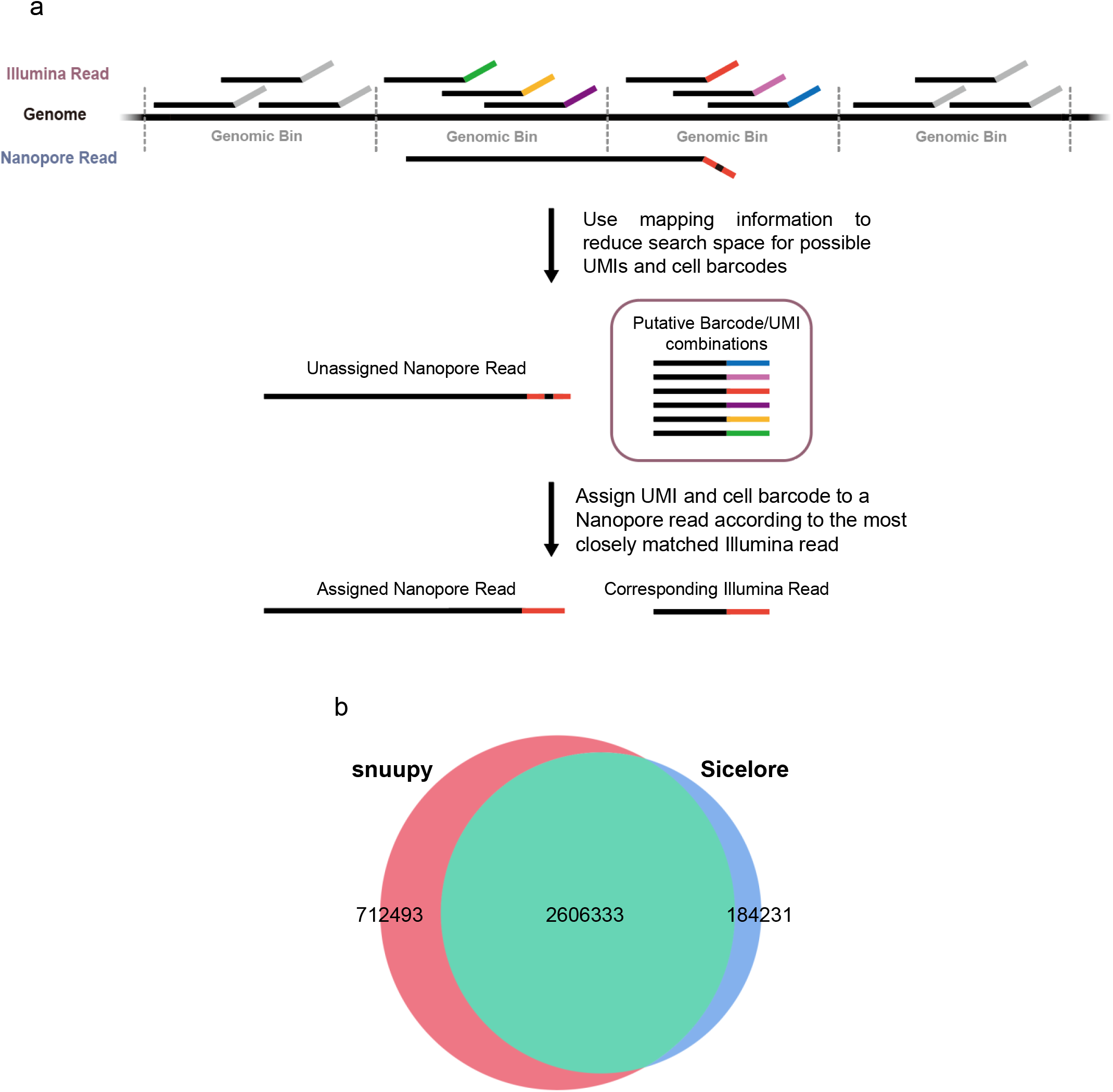
Snuupy assigns cell barcodes and UMIs for Nanopore reads according to the information from Illumina data. **a**, Snuupy uses mapping information to reduce the search space as previously reported in Sicelore. **b**, Overlap between snuupy and Sicelore allocated reads.

**Supplemental Figure 9.**
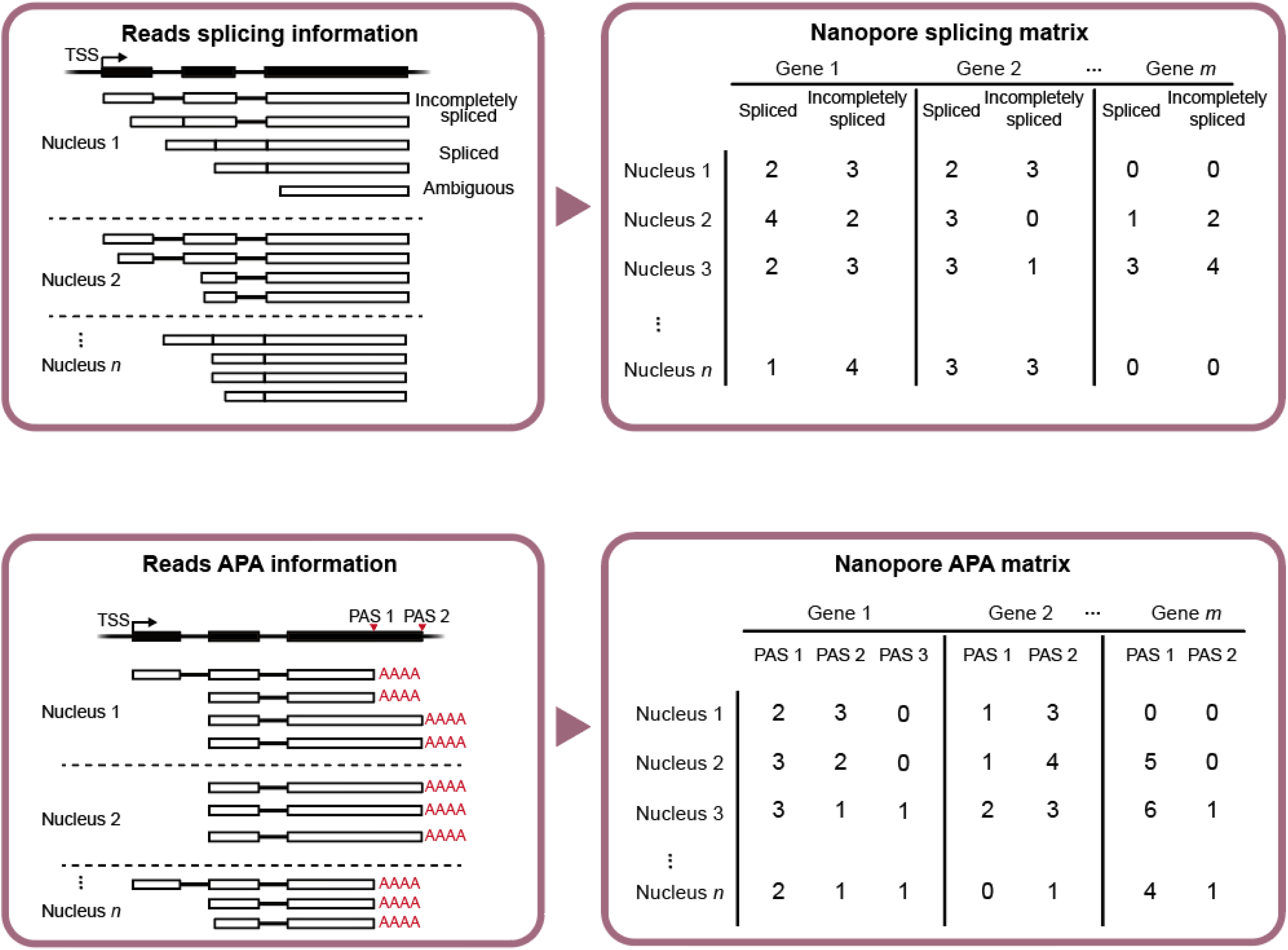
Scheme for deriving the splicing and APA matrices from Nanopore data.

